# An ABBA-BABA Test for Introgression Using Retroposon Insertion Data

**DOI:** 10.1101/709477

**Authors:** Mark S. Springer, John Gatesy

## Abstract

DNA sequence alignments provide the majority of data for inferring phylogenetic relationships with both concatenation and coalescence methods. However, DNA sequences are susceptible to extensive homoplasy, especially for deep divergences in the Tree of Life. Retroposon insertions have emerged as a powerful alternative to sequences for deciphering evolutionary relationships because these data are nearly homoplasy-free. In addition, retroposon insertions satisfy the ‘no intralocus recombination’ assumption of summary coalescence methods because they are singular events and better approximate neutrality relative to DNA sequences commonly applied in phylogenomic work. Retroposons have traditionally been analyzed with phylogenetic methods that ignore incomplete lineage sorting (ILS). Here, we analyze three retroposon data sets for mammals (Placentalia, Laurasiatheria, Balaenopteroidea) with two different ILS-aware methods. The first approach constructs a species tree from retroposon bipartitions with ASTRAL, and the second is a modification of SVD-Quartets. We also develop a χ^2^ Quartet-Asymmetry Test to detect hybridization using retroposon data. Both coalescence methods recovered the same topology for each of the three data sets. The ASTRAL species tree for Laurasiatheria has consecutive short branch lengths that are consistent with an anomaly zone situation. For the Balaenopteroidea data set, which includes rorquals (Balaenopteridae) and gray whale (Eschrichtiidae), both coalescence methods recovered a topology that supports the paraphyly of Balaenopteridae. Application of the χ^2^ Quartet-Asymmetry Test to this data set detected 16 different quartets of species for which historical hybridization may be inferred, but significant asymmetry was not detected in the placental root and Laurasiatheria analyses.

## Introduction

Retroelements are a broad class of ‘copy-and-paste’ transposable elements that comprise ~90% of the three million transposable elements in the human genome (Bannert and Kruth, 2004). The two major groups of retroelements are defined by the presence/absence of long-terminal repeats (LTRs). Non-LTR retroelements include short interspersed nuclear elements (SINEs) and long interspersed nuclear elements (LINEs), which are commonly referred to as retroposons (Churakov et al. 2009; Nishihara et al. 2009). LTR retroelements include LTR-retrotransposons and endogenous retroviruses.

In the last two decades, retroelement insertions have emerged as powerful tools for resolving phylogenetic relationships (Shimamura et al. 1997; Shedlock and Okada 2000; Nikaido et al. 2001; Ray et al. 2006; Nilsson et al. 2010, 2012; Hartig et al. 2013; Doronina et al. 2015, 2017a,b). In part, this is because retroelement insertions have exceptionally low levels of homoplasy relative to sequence-based analyses (Shedlock et al. 2000; Shedlock et al. 2004; Hallström et al. 2011; Kuritzin et al. 2016; Doronina et al. 2017b, 2019; Gatesy et al. 2017). Conflicting retroposons are generally the products of incomplete lineage sorting (ILS) rather than homoplasy (Robinson et al. 2008; Suh et al. 2011; Kuritzin et al. 2016). Avise and Robinson (2008) suggested the term hemiplasy for outcomes of ILS that mimic homoplasy. Indeed, the basic mechanics of the cut and paste process that generates retroposon insertions across the genome contrasts with the process that yields high homoplasy in DNA sequence data. In the latter, multiple substitutional mutations at single sites commonly flip to and fro between just four states (G, A, T, C).

Retroposon insertions have commonly been analyzed using one or more variants of parsimony (e.g., Camin-Sokal parsimony [Nikaido et al. 1999]; polymorphism parsimony [Suh et al. 2015]; Dollo parsimony [Lammers et al. 2019]), distance methods (e.g., neighbor joining [Lammers et al. 2019]), or networks (e.g., Doronina et al, 2017a; Lammers et al. 2019). More recently, Springer and Gatesy (2019) suggested that retroposons have several properties that are well matched to the assumptions of summary coalescent methods such as ASTRAL (Mirarab and Warnow 2015). First, summary coalescent methods assume that ILS is the only source of gene tree heterogeneity. However, gene-tree reconstruction error is a major source of gene-tree heterogeneity for summary coalescent analyses with sequence-based gene trees (Huang et al. 2010; Patel et al. 2013; Gatesy and Springer 2014; Springer and Gatesy 2016; Scornavacca and Galtier 2017). By contrast, very low levels of homoplasy effectively liberate retroposon insertions from gene tree reconstruction error. Second, the basic units of analysis in a summary coalescent analysis are coalescence genes (c-genes), segments of the genome for which there has been no recombination over the phylogenetic history of a clade (Doyle, 1992, 1997). Summary coalescent methods assume that there has been free recombination between c-genes but no recombination within c-genes. This assumption is routinely violated when complete protein-coding sequences are employed as c-genes because individual exons are often stitched together from far-flung regions of the genome (Gatesy and Springer, 2013; Springer and Gatesy, 2016, 2018; Mendes et al. 2019). C-genes become even shorter as more taxa are added to an analysis because of the ‘recombination ratchet’ (Gatesy and Springer 2014; Springer and Gatesy 2016, 2018). By contrast, the presence/absence of a retroposon is almost always due to a single mutational event (Doronina et al. 2019) and is not subject to intralocus recombination. The singularity of each retroposon insertion nullifies problems derived from the recombination ratchet. Third, retroposons are largely neutral markers (Kuritzin et al. 2016; Chuong et al. 2017; Kubiak and Makalowska 2017; Doronina et al. 2019) as is assumed by summary coalescent methods (Liu et al. 2009). More specifically, the overwhelming majority of retroposon insertions occur in genomic ‘safe havens’ where they have no known functional or selective significance (Kuritzin et al. 2016; Chuong et al. 2017). In addition, single-site coalescent methods such as SVDquartets (Chifman and Kubatko 2014) and SVDquest (Vachaspata and Warnow 2018) can be used to estimate a species tree from unlinked SNP data. These methods bypass the problem of gene tree estimation error that afflicts summary coalescent analyses with sequence-based gene trees (Chou et al. 2015), but the assumption of free recombination among sites is violated when SNP methods are applied to linked sites in a gene. Individual retroposon insertions typically occur at unlinked sites and are more appropriate than gene sequences for analysis with single-site coalescent methods. Finally, as is the case for any new mutation, a retroposon insertion is polymorphic when it first appears in a population (e.g., Lammers et al. 2019) as is required for any marker that is subject to ILS.

Given that the properties of retroposon insertions are well matched to the assumptions of coalescent methods for species tree estimation, the distribution of conflicting retroposons should fit the predictions of ILS under random mating unless there is also introgression or lateral transfer of genomic material (Kuritzin et al. 2016). Kuritzin et al. (2016) developed an approach for inferring introgression based on the distribution of conflicting retroposons among three taxa relative to an outgroup that lacks the insertions. This framework, named KKSC after the last names of the four authors, does not consider cases where a retroposon insertion originates in the common ancestor of all four lineages and then undergoes ILS. Importantly, this scenario will result in both 0011 and 1100 patterns that are both consistent with the quartet split AA|BB (Fig. 1A,B). Kuritzin et al.’s (2016) method only tallies the 0011 pattern (Fig. 1B). Nevertheless, both 0011 and 1100 patterns are expected under the MSC with ILS. Such patterns are routinely analyzed with ABBA-BABA tests for DNA sequence data (Green et al. 2010).

**Fig. 1.**
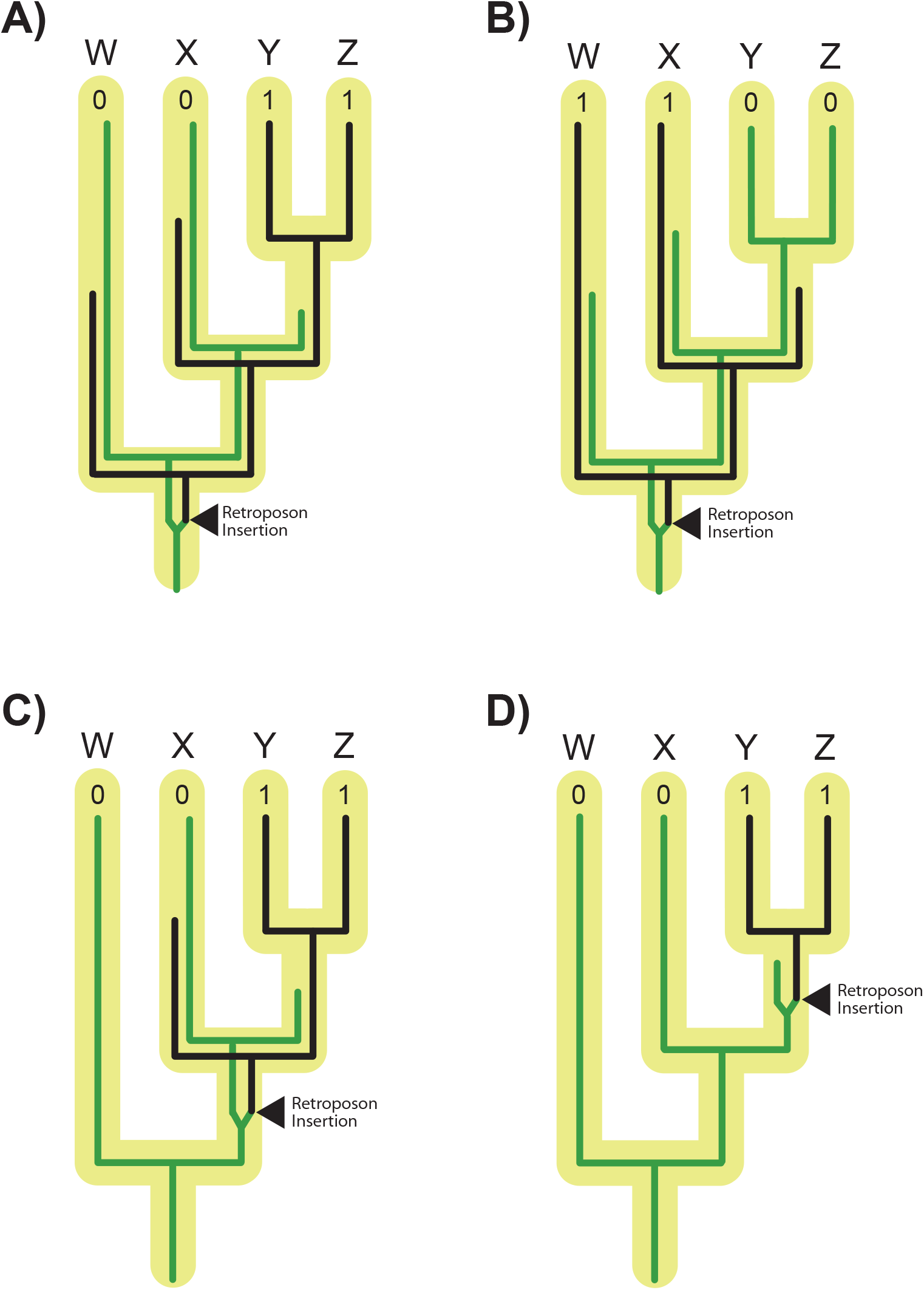
Schematic of different scenarios that will result in 0011 and 1100 retroposon insertion patterns. These scenarios allow for synapomorphic changes and hemiplasy, but not homoplasy. In (A), a retroposon insertion originates in the common ancestor of all four lineages (W, X, Y, Z) and results in a 0011 pattern through ILS. In (B), a retroposon originates in the common ancestor of all four lineages and results in a 1100 pattern through ILS. In (C), a retroposon insertion that originates in the common ancestor of just three lineages (X, Y, Z) will sometimes support 0011 even though the insertion alleles in Y and Z do not coalesce in the most recent common ancestor of Y and Z. In (D), a synapomorphic retroposon originates and is fixed in the common ancestor of Y and Z, resulting in a 0011 pattern.

Here, we construct species trees for three retroposon data sets (Nishihara et al. 2009; Doronina et al. 2017a; Lammers et al. 2019) using two coalescent methods that account for ILS. Then, we develop a general χ^2^ ABBA-BABA test (Green et al. 2010) for retroposon insertions. This test allows for insertions that originate in the common ancestor of the entire clade as well as on more crownward branches (Fig. 1A-D), and provides a statistical framework to determine if ILS alone or ILS + introgression are required to explain the distribution of conflicting retroposons. Because retroposon insertions are low-homoplasy characters, even for species trees with relatively ancient divergences and short internodes, we argue for the utility of our approach for discriminating ILS versus hybridization among lineages that diverged as deep as the upper Cretaceous. Analogous ABBA-BABA tests using sequence-based gene trees at this depth are, by contrast, challenged by multiple overlapping substitutions, model-mis-specification, and poorly reconstructed gene trees.

## Methods

### Retroposon Data Sets

The first data set is comprised of 68 LINE insertions that are informative for resolving relationships among three subclades of Placentalia (Afrotheria [e.g., elephants, hyraxes], Boreoeutheria [e.g., primates, rodents, bats], Xenarthra [e.g., sloths, armadillos]) that diverged from each other at the base of Placentalia (Nishihara et al. 2009). Retroposon counts for the three different resolutions of this clade are 22 for Boreoeutheria + Xenarthra, 25 for Afrotheria + Boreoeutheria, and 21 for Afrotheria + Xenarthra. Each taxon in the dataset is coded as absent (0) or present (1) for each retroposon character (i.e., there are no missing data). Previously, Nishihara et al. (2009) concluded that these three lineages diverged from each other nearly simultaneously in the Cretaceous.

The second data set is comprised of 102 retroposons (76 LINEs, 23 LTRs, and three retropseudogenes) for four orders of Laurasiatheria (Carnivora [e.g., dogs, cats], Cetartiodactyla [e.g., pigs, cows, whales], Chiroptera [bats], Perissodactyla [e.g., horses, rhinos]) and the outgroup Eulipotyphla (e.g., shrews, moles, hedgehogs) (Doronina et al. 2017a). For this data set, retroposon insertions support ten different bipartitions that are phylogenetically informative for resolving ingroup relationships. Retroposon counts for the Laurasiatheria data set range from four to 14 for different bipartitions and were taken from figure 1 in Doronina et al. (2017a). There are no missing data for these 102 retroposon insertions. We also analyzed an expanded data set of 162 retroposons that included 60 additional loci, of which 16 were scored as missing for the eulipotyphlan representative and 16 were scored as present in Eulipotyphla and a subset of other laurasiatherians (supplementary table 1 of Doronina et al. 2017a).

The third retroposon data set is based on a genome-wide scan that identified 24598 CHR-SINE insertions for balaenopteroid baleen whales (Lammers et al. 2019). These SINE insertions occur in two or more of six balaenopteroid species, but not in the outgroup Balaenidae. The six balaenopteroids include five balaenopterids (*Balaenoptera acutorostrata* [common minke whale], *B. musculus* [blue whale], *B. borealis* [sei whale], *B. physalus* [fin whale], *Megaptera novaeangliae* [humpback whale]) and one eschrichtiid (*Eschrichtius robustus* [gray whale]). The balaenid outgroup is *Eubalaena glacialis* [North Atlantic right whale]). This data set consisted of 57 different bipartitions with counts that ranged from 46 to 7373 (Table S1). There are no missing data in the balaenopteroid dataset.

### Species Tree Estimation Using Retroposon Insertions

We estimated species trees using two ILS-aware methods (Fig. 2). First, we employed the protocol outlined previously by Springer and Gatesy (2019). Each retroposon matrix of presence/absence characters was converted into a set of incompletely resolved ‘gene trees’ with each gene tree representing a single retroposon character and supporting just a single bipartition. Summary coalescent analysis of each data set was then executed using ASTRAL-III v. 5.6.1 (Zhang et al. 2017), which accepts input of incompletely resolved gene trees (Fig. 2A). The output of ASTRAL includes the inferred species tree, estimated branch lengths in coalescent units (CUs), and local posterior probabilities (PPs) for each supported clade (Sayyari and Mirarab 2016). ‘Gene-wise’ bootstrapping in ASTRAL gave a second measure of clade support. Simmons et al. (2019) recently argued that genewise resampling is more appropriate than site-wise bootstrap resampling, but for gene trees that represent single retroposon insertions, these two support measures are essentially identical. Finally, we tested the stability of supported clades in a third way by calculating the minimum number of gene tree removals that are required to collapse each clade. Specifically, the protocol outlined by Gatesy et al. (2019) was implemented, and this procedure estimates the number of gene tree removals necessary to yield the best ASTRAL tree that lacks the clade of interest.

**Fig. 2.**
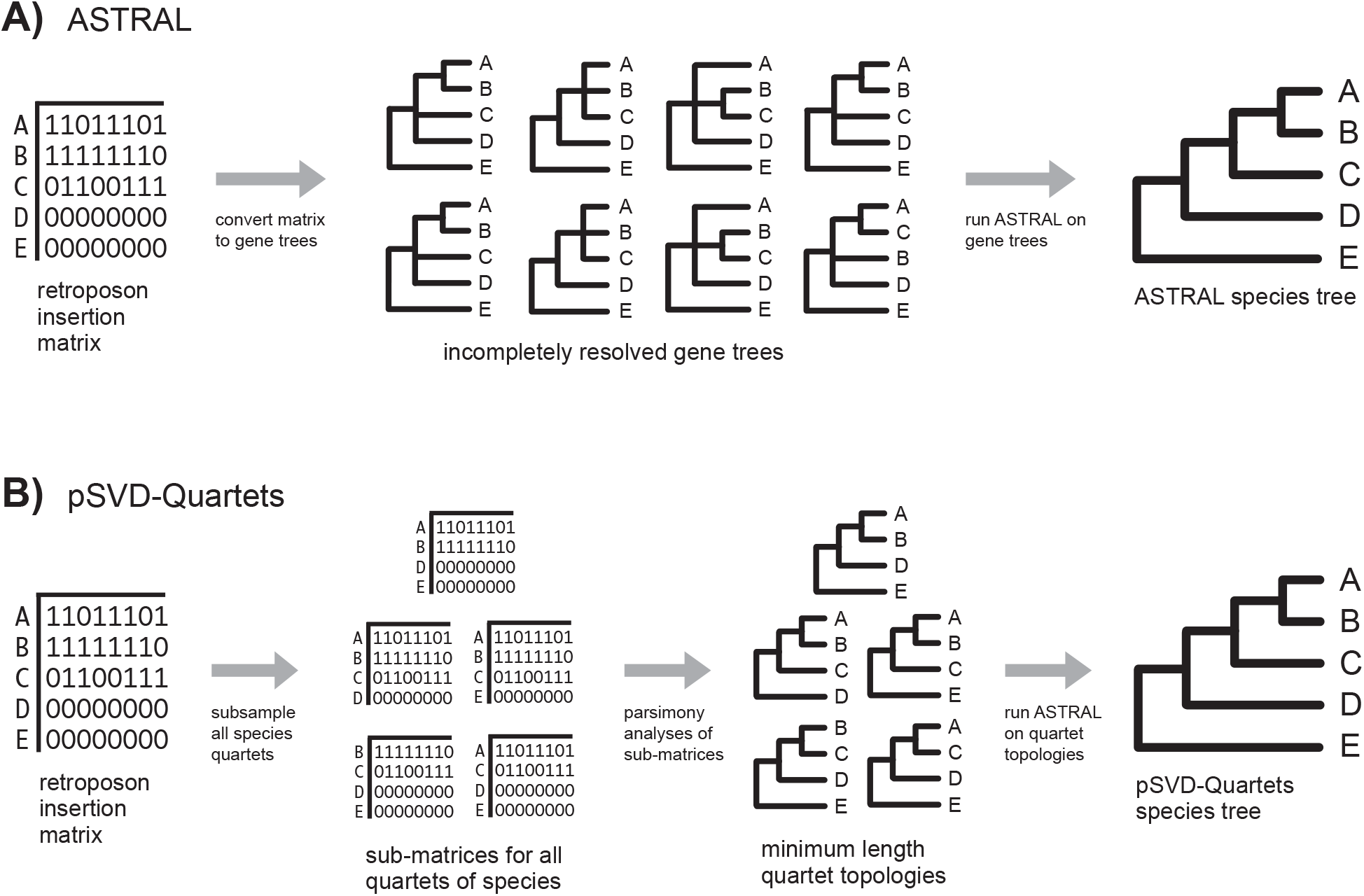
Schematic of two coalescence methods that can be used to reconstruct a species tree from a retroposon insertion matrix. In (A), ASTRAL is used to analyze partially resolved gene trees (Springer and Gatesy 2019). In (B), retroposon insertions are analyzed with pSVD-Quartets. Taxon E is the specified outgroup in this hypothetical example.

We also analyzed each retroposon matrix using a variant of the summary coalescence method SVDquartets (Chifman and Kubatko 2014) to confirm species-tree topologies based on ASTRAL analysis. SVDquartets sensu stricto (Chifman and Kubatko 2014) originally was implemented to analyze nucleotide SNPs with a general time-reversible (GTR) model or Jukes-Cantor (JC) model of DNA sequence evolution. For retroposon insertions, we developed a variant of SVDquartets that we call ‘Parsimony SVD-Quartets’ (pSVD-Quartets). This method utilizes (1) parsimony analyses of all possible species quartets and (2) subsequent assembly of optimal parsimony resolutions for each quartet of species to build a species tree (Fig. 2B). To implement pSVD-Quartets, we used the “GetTrees” command in PAUP* 4.0a (build 165) (Swofford, 2002) to import each dataset of retroposon gene trees (newick format) that we had assembled for the ASTRAL analyses above. Next, we used the MatrixRep command to save the MRP matrix of nodal characters. This MRP matrix was then executed, and the informative characters were exported into a new nexus file, which was then used to determine the optimal tree for each quartet of taxa. The first data set (Placentalia) included a single quartet with one optimal solution. The Laurasiatheria data set, in turn, included five taxa, five quartets and six optimal trees. The additional tree for Laurasiatheria resulted from a quartet with two equally parsimonious trees. Here, we downweighted the two trees by 0.5X so that each of the five quartets was given the same weight in parsimony analysis. Finally, for the balaenopteroid data set with seven taxa there were 35 quartets and 35 optimal trees. After determining the optimal parsimony tree(s) for each quartet of taxa, we then used ASTRAL to determine the species tree based on the full set of optimal quartet trees for each data set (Fig. 2B). In a similar procedure, SVDquest (Vachaspati and Warnow 2018) utilizes ASTRAL to find the optimal species tree from a set of quartet trees. We used the exact search option in ASTRAL to search for the species tree topology that fits the most input quartets trees. The resulting branch lengths and local PP support scores in the ASTRAL output have no coherent meaning for pSVD-Quartets topologies, because each of the input quartets is based on parsimony analysis of multiple unlinked loci instead of individual unlinked loci, so these values were ignored.

### Test for Introgression

For each data set (Placentalia, Laurasiatheria, Balaenopteroidea), we tested whether the conflicts among retroposon insertions could be explained by ILS alone. The χ^2^ Quartet-Asymmetry Test is a simple χ^2^ variant of the ABBA-BABA test (Green et al. 2010) that can be applied to both symmetric and pectinate four-taxon subtrees (Fig. 3). The rationale for application of this test to symmetric subtrees is that each of the two alternative and symmetric quartet trees are equally likely under the MSC, as is the case for the ABBA-BABA test with pectinate trees (Fig. 4). This is because the alternate quartet resolutions have equal opportunities (branch lengths) for ABBA (0110 or 1001) and BABA (1010 or 0101) patterns under ILS (Fig. 4). For each ASTRAL species tree (Placentalia, Laurasiatheria, Balaenopteroidea), we examined each unique quartet of species and compared the observed and expected number of retroposons that support each of the two quartet trees that conflict with the inferred species tree. For each data set, we imported all of the incompletely resolved gene trees into PAUP* 4.0a165 (Swofford 2002); each gene tree corresponds to a single retroposon insertion (see Fig. 2A). Next, we pruned all taxa from these gene trees except for the four taxa that comprise the relevant quartet. Gene trees that were missing one or more of the four taxa were also discarded. We then computed the majority rule consensus of these trees for each quartet with the option to show the frequencies of all observed bipartitions. Finally, we used this output to determine the numbers of gene trees (i.e., retroposon insertions) that support the quartet in the species tree and each of the alternative conflicting quartet trees. The expectation under ILS without hybridization is that the number of retroposon characters that support each of the two non-species-tree quartets will be equal (Degnan and Rosenberg 2009; Ané 2010). By contrast, introgression has the potential to distort this 50:50 ratio, with one alternative quartet tree overrepresented relative to the alternative quartet tree. Because multiple quartets of species were examined in each species tree, we employed a Holm-Bonferroni sequential correction (Holm 1979) with an EXCEL calculator (Gaetano 2013) to account for multiple comparisons.

**Fig. 3.**
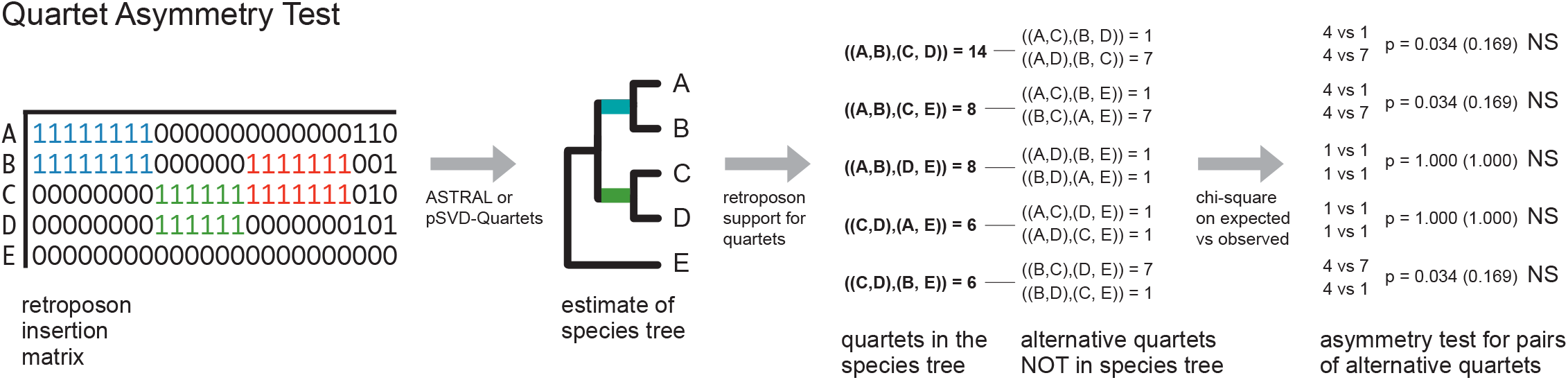
Schematic showing χ^2^ Quartet-Asymmetry Tests for a retroposon matrix. The first step is to construct a species tree using ASTRAL or pSVD-Quartets (Fig. 2). Next, each combination of four species is examined individually to determine if the two alternative quartets that are not in the species tree are equally supported by retroposon insertions as expected under ILS. A χ^2^ test with a correction for multiple tests is used to assess statistical significance. Quartets of species with significant differences in the number of alternative quartets provide evidence for introgression pathways. Note that in this hypothetical case, three quartets show significant asymmetery before, but not after, correction for multiple tests. Retroposons that support clade A+B are colored blue, retroposons that support clade C+D are colored green, and strong conflict for B+C is red.

**Fig. 4.**
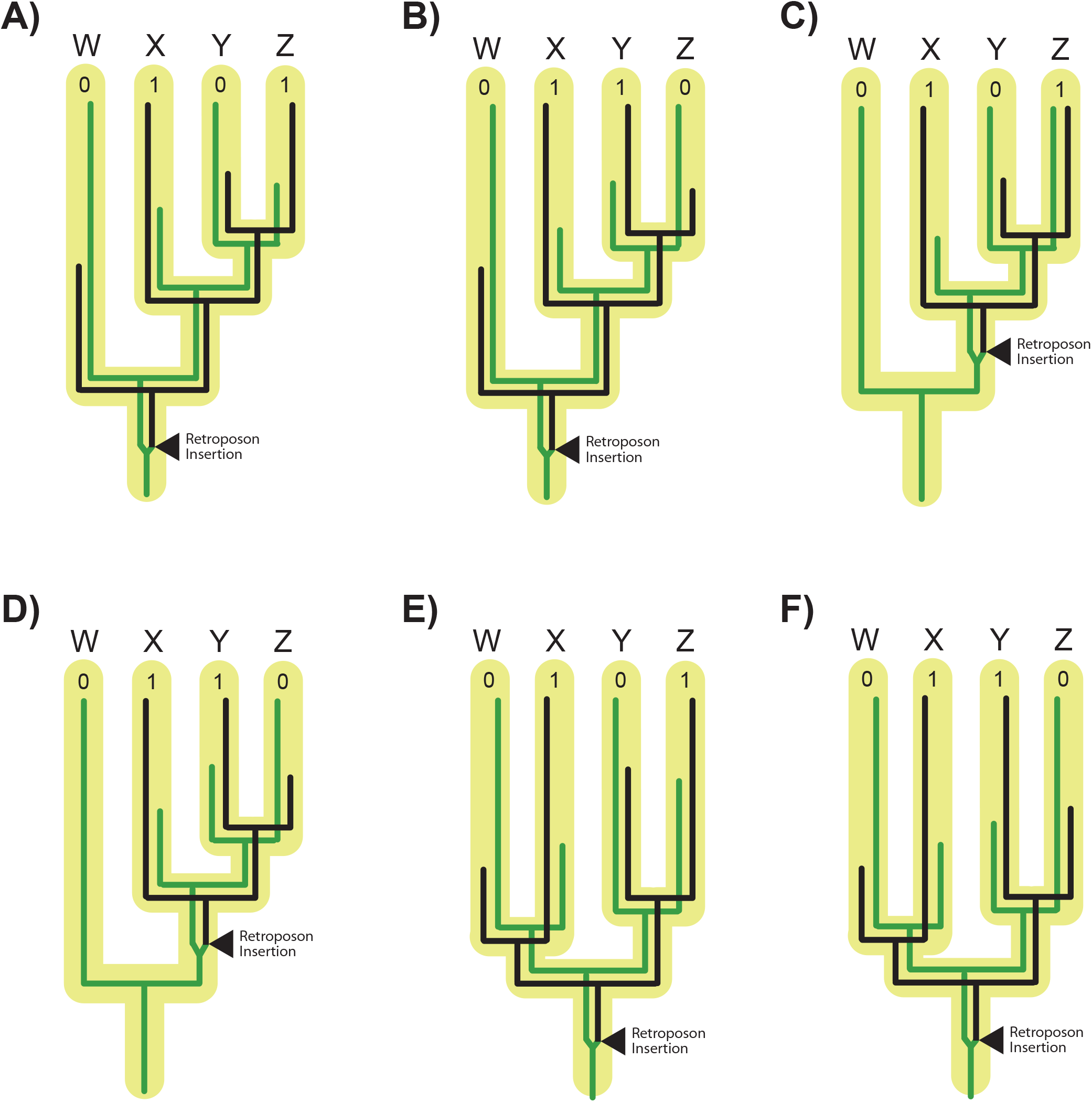
Illustration of ABBA-BABA patterns for the distribution of retroposon insertions on pectinate and symmetric trees for four-taxa (W, X, Y, Z). In all cases, the quartet split on the unrooted species tree is WX|YZ. In (A-D), pectinate trees have equivalent probabilities for ABBA and BABA hemiplasy via ILS under neutrality. ABBA (0110 or 1001) and BABA (0101 or 1010) patterns can arise if the retroposon insertion originates in the ancestor of all four taxa (A-B) or in the ancestor of X, Y, and Z (C-D). Panels A and B show equivalent opportunities for 0101 and 0110 patterns, but similar opportunities exist for 1010 and 1001 patterns. In (E-F), symmetric trees have equivalent opportunities for ABBA and BABA hemiplasy with ILS under neutrality, but will only occur when the retroposon insertion originates in the ancestor of all four taxa.

### Simulation of Gene Trees

Ten thousand gene trees were simulated from the ASTRAL retroposon species tree (102 retroposons) for Laurasiatheria (branch lengths in CUs) using DendroPy v3.12.0 (Sukumaran and Holder 2010) and a script from Mirarab et al. (2014). In anomaly zone conditions, the most probable gene tree does not match the species tree (Degnan and Rosenberg 2006, 2009). The simulated gene tree output was used to determine whether the most common gene tree topology contradicts the species tree.

## Results and Discussion

### Placentalia Retroposons

The ASTRAL and pSVD-Quartets species trees for the placental root data set of 68 retroposon insertions both resolve Afrotheria + Boreoeutheria, but with low bootstrap support (58%) and local PP (0.52) values on the ASTRAL tree (Fig. 5A). Just three gene tree removals collapse the clade, further suggesting instability and low support due to extensive conflicts in the retroposon insertion data. The internal branch length uniting Afrotheria and Boreoeutheria is very short, just 0.0445 CUs, especially relative to long terminal branches that extend to the Cretaceous (Fig. 5B; Meredith et al. 2011). Kuritzin et al. (2016) performed their KKSC test on this same data set and concluded that there is no compelling evidence for ancient hybridization among lineages. In agreement with Kuritzin et al. (2016), we did not find any support for introgression based on the single species quartet (outgroup, Xenarthra, Afrotheria, Boreoeutheria) that we examined with the χ^2^ Quartet-Asymmetry Test (p = 0.91). The mixed support that retroposons provide for different resolutions of the placental root is consistent with the conflicting results of sequence-based phylogenetic studies that favor Xenarthra + Boreoeutheria (Murphy et al. 2001; Meredith et al. 2011 [DNA], McCormack et al. 2012; Romiguier et al. 2013), Afrotheria + Boreoeutheria (Waddell et al. 2001), and Afrotheria + Xenarthra (Song et al. 2012; Meredith et al. 2011 [amino acids], Morgan et al. 2013; Tarver et al. 2016; Chen et al. 2017; Liu et al. 2017).

**Fig. 5.**
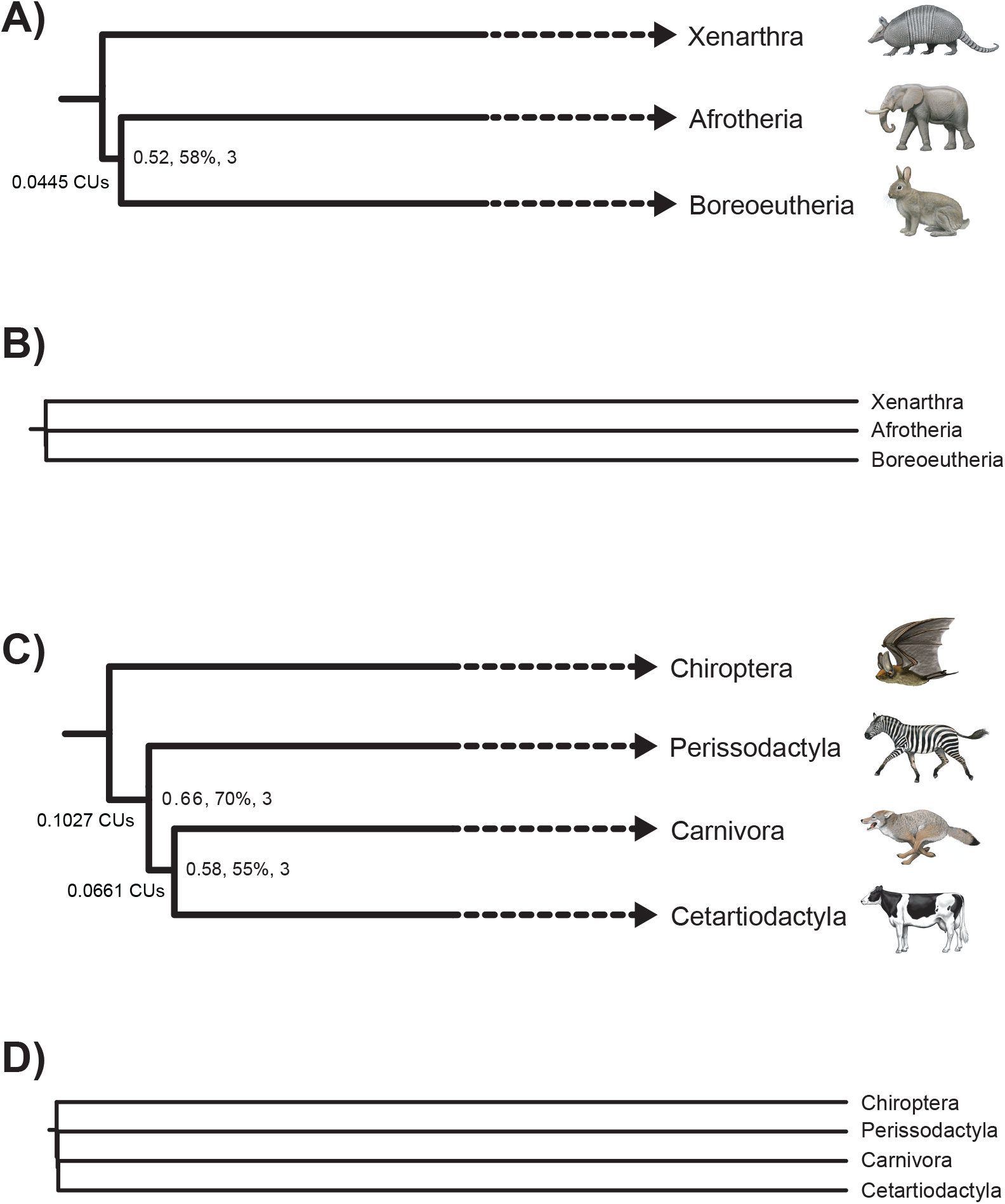
ASTRAL species trees for Placentalia (A-B) and Laurasiatheria (C-D). Numbers at internal nodes in (A) and (C) correspond to local PP, bootstrap support percentage, and the minimum number of gene trees that must be removed to collapse the clade. Terminal branches are truncated in (A) and (C) so that relative lengths of internal branches in coalescent units (CUs) can be seen. Species trees in (B) and (C) also show branch lengths in CUs; CU estimates for terminal branches are derived from molecular clock analyses (Meredith et al. 2011) because ASTRAL does not estimate terminal branch lengths. One million years from timetree estimates were converted to one CU (see Springer and Gatesy 2016 for rationale). Relationships in pSVD-Quartets species trees are identical to those in the ASTRAL species trees that are shown. The successive short branch lengths of 0.1027 and 0.0661 CUs in the species tree for Laurasiatheria (C-D) are consistent with anomaly zone conditions (Degnan and Rosenberg 2006).

### Laurasiatheria Retroposons

We re-examined the Laurasiatheria polytomy using 102 retroposon insertions from Doronina et al. (2017a). This polytomy is perhaps the most challenging interordinal problem in all of Placentalia given that it involves four lineages (Carnivora + Pholidota, Cetartiodactyla, Chiroptera, Perissodactyla) that diverged in the late Cretaceous and two very short internal branches (Fig. 5D; Meredith et al. 2011; dos Reis et al. 2012; Emerling et al. 2015; Foley et al. 2016; Tarver et al. 2016). Previous investigations of this systematic problem have employed both genome-scale DNA sequence data and retroposon insertions, yielding highly incongruent results (Nishihara et al. 2006; Hallström et al. 2011; Meredith et al. 2011; Song et al. 2012; Tarver et al. 2016; Chen et al. 2017; Doronina et al. 2017a; Feijoo and Parada, 2017; Liu et al. 2017). Some of these analyses are further complicated by homology problems with the underlying data (see Springer and Gatesy 2016, 2017; Gatesy and Springer 2017). Our ASTRAL and pSVD-Quartets analyses of Doronina et al.’s (2017a) data set recovered the same species tree reported by these authors based on polymorphism parsimony (Felsenstein 1979). Carnivora and Cetartiodactyla are sister taxa, with Perissodactyla and Chiroptera as successively more distant relatives to this clade. However, both posterior probabilities and bootstrap scores for interordinal relationships are low, and each clade collapses with the removal of just three gene trees (Fig. 5C).

Branch lengths for the consecutive internal branches at the base of the tree are very short (0.1027, 0.0661). These tiny branch lengths, along with the pectinate species tree, suggest that the virtual polytomy for Laurasiatheria may reside in the anomaly zone (Degnan and Rosenberg 2006, 2009) wherein the most likely gene tree topology differs from the species tree. Consistent with this interpretation, we simulated 10000 genes trees from the ASTRAL species tree, and the three most common gene tree topologies differ from the species tree. Specifically, all three of the symmetrical gene trees for the ingroup are better represented than the pectinate gene tree that agrees with the species tree. The symmetrical gene trees occur with frequencies of 1182 ((Cetartiodactyla, Carnivora),(Perissodactyla, Chiroptera)), 1046 (Cetartiodactyla, Perissodactyla), (Carnivora, Chiroptera)), and 991 ((Cetartiodactyla, Chiroptera),(Carnivora, Perissodactyla)). By contrast, there are only 916 gene trees that agree with the species tree (Chiroptera,(Perissodactyla,(Carnivora, Cetartiodactyla))). An anomaly zone situation has also been suggested for palaeognath birds (Cloutier et al. 2019; Sackton et al. 2019), but in this case, the consecutive short branches are more likely due to conflicts from high reconstruction error for sequence-based gene trees, an interpretation that is corroborated by analysis of retroposon insertion data from these birds (Springer and Gatesy 2019).

Doronina et al. (2017a) extended the KKSC test of Kuritzin et al. (2016) to a 4-Lineage Insertion Likelihood test that allowed for different hybridization scenarios. They concluded that gene flow among lineages had occurred and that the most likely hybridization pathways were between (1) Perissodactyla and Chiroptera, (2) Perissodactyla and Carnivora, and (3) Perissodactyla and the common ancestor of Carnivora + Cetartiodactyla. We applied the χ^2^ Quartet-Asymmetry Test to all five species quartets for this data set and found that there are no statistically significant differences between the observed and expected numbers of retroposons for pairs of non-species-tree quartets. This pattern is consistent with ILS and does not require introgression (Table 1). We also analyzed Doronina et al.’s (2017a) expanded data set with 162 retroposon insertions, some of which were scored as missing or present for the eulipotyphlan outgroup, and again obtained non-significant results for the χ^2^ Quartet-Asymmetry Test (Table 1). The most extreme discrepancy in these comparisons is 20 versus 12 retroposon insertions for alternative non-species tree quartets, but the p value (0.157) for this comparison is not significant even without correcting for multiple comparisons. This disagrees with Doronina et al.’s (2017a) conclusion that there is evidence for hybridization among basal laurasiatherian lineages based on their analysis of the data set with 162 retroposon insertions. Either the χ^2^ Quartet-Asymmetry Test is less powerful than Doronina et al.’s (2017a) hybridization test or their inferences are overconfident.

**Table 1.** The number of retroposons that support different splits for five different quartets of taxa that are associated with the Laurasiatheria polytomy^a^.

| Quartet of taxa | Split | 102 Retroposon Data set |  | 162 Retroposon Data Set<sup>b</sup> |  |
| --- | --- | --- | --- | --- | --- |
|  |  | Obs. #.<br>quartets | Exp. #<br>quartets under<br>ILS | Obs. #.<br>quartets | Exp. #<br>quartets under<br>ILS |
| Eu, Pe, Ca, Ce | **Eu Pe \| Ca Ce** | **22** |  | **25** |  |
|  | Eu Ca \| Pe Ce | 22 | 21 | 22 | 21.5 |
|  | Eu Ce \| Pe Ca | 20 | 21 | 21 | 21.5 |
| Eu, Pe, Ca, Ch | **Eu Ch \| Ca Pe** | **25** |  | **25** |  |
|  | Eu Ca \| Pe Ch | 23 | 20 | 23 | 20 |
|  | Eu Pe \| Ca Ch | 17 | 20 | 17 | 20 |
| Eu, Pe, Ce, Ch | **Eu Ch \| Pe Ce** | **24** |  | **24** |  |
|  | Eu Ce \| Pe Ch | 20 | 16 | 21 | 17.5 |
|  | Eu Pe \| Ce Ch | 12 | 16 | 14 | 17.5 |
| Eu, Ca, Ce, Ch | **Eu Ch \| Ca Ce** | **28** |  | **31** |  |
|  | Eu Ce \| Ca Ch | 18 | 17 | 19 | 18.5 |
|  | Eu Ca \| Ce Ch | 16 | 17 | 18 | 18.5 |
| Pe, Ch, Ce, Ca | **Pe Ch \| Ce Ca** | **25** |  | **38** |  |
|  | Ch Ce \| Pe Ca | 15 | 17 | 25 | 24 |
|  | Ch Ca \| Pe Ce | 19 | 17 | 23 | 24 |

<sup>a</sup>The species tree split for each quartet is shown in bold font.
<sup>b</sup>Compiled from Doronina et al.'s (2017a) Supplemental Table S1.
Abbreviations: Ca, Carnivora; Ce, Cetartiodactyla; Ch, Chiroptera; Eu, Eulipotyphla; Pe, Perissodactyla.

### Balaenopteroidea Retroposons

Phylogenetic relationships among balaenopteroid species remain contentious based on previous phylogenetic analyses. Some studies support a sister-group relationship between Eschrichtiidae and Balaenopteridae (e.g., Deméré et al. 2008; Steeman et al. 2009; Marx 2010; Gatesy et al. 2013), whereas Eschrichtiidae is nested inside of a paraphyletic Balaenopteridae in other work (e.g., McGowen et al. 2009; Marx and Fordyce 2015). Figure 6A,B shows the ASTRAL species tree for balaenopteroids with internal branch lengths in CUs. Relationships in the pSVD-Quartets topology (not shown) are identical to those in the ASTRAL tree. The basal split among balaenopteroids is between minke whale and the other five balaenopteroids. Consistent with previous molecular studies (e.g., Rychel et al. 2004; Sasaki et al. 2005; Nikaido et al. 2006; Deméré et al. 2008; McGowen et al. 2009; Árnason et al. 2018), fin whale and humpback whale are sister taxa, and blue whale is closely related to sei whale. The gray whale is positioned as the sister taxon to fin whale + humpback whale. This coalescence-based retroposon tree matches a neighbor-joining distance analysis and a Bayesian inference tree for the same data (Lammers et al. 2019) as well as earlier molecular and total evidence work (McGowen et al. 2009; Marx and Fordyce, 2015; Árnason et al. 2018). The branch that unites gray whale with the fin whale + humpback whale clade is very short (0.0182 CUs) on the ASTRAL tree, although there is moderately high bootstrap support (89%) and local PP (0.88) for this clade which is resistant to collapse following removal of many gene trees from ASTRAL analysis. The nested position of gray whale inside of Balaenopteridae renders this family paraphyletic (Fig. 6A).

**Fig. 6.**
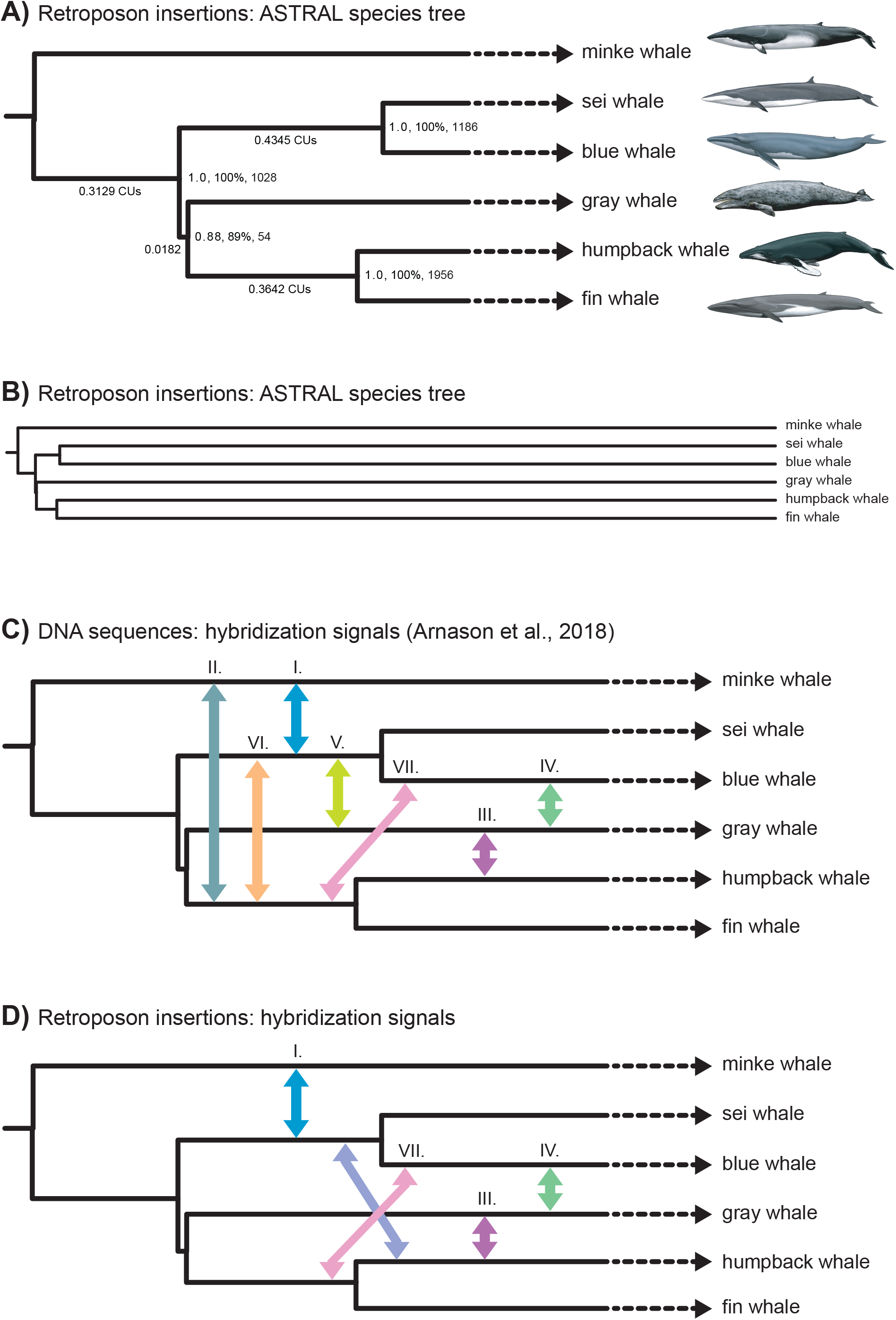
The ASTRAL species tree for Balaenopteroidea (A) based on 24598 retroposon bipartitions (A-D). Numbers at internal nodes correspond to local PP, bootstrap support percentage, and the minimum number of gene trees that must be removed to collapse the clade. Terminal branches are truncated in (A), (C), and (D) so that relative lengths of internal branches in coalescent units (CUs) can be seen. The species trees in (B) also shows estimates of terminal branches in CUs that were derived from a molecular clock analysis (McGowen et al. 2009). One million years was converted to one CU (see Springer and Gatesy, 2016). Relationships in the pSVD-Quartets species tree are identical to those in the ASTRAL species tree that is shown. In (C), Árnason et al.’s (2018) seven hypothesized pathways for introgression that are based on genomic sequences are shown. Possible gene flow pathways detected using the χ^2^ Quartet-Asymmetry Test for retroposon insertions are shown in (D).

Overall, our species tree is robustly supported at most nodes and concordant with the ASTRAL tree that Árnason et al. (2018) reported for the same taxa based on gene trees for 34192 genome fragments, each of which was 20 kb in length. Unlike retroposon gene trees, there are significant non-biological contributors to gene tree heterogeneity when gene trees are inferred from DNA sequence alignments. As noted above, these include long-branch misplacement, misaligned sequences, long tracts of missing data for one or more species, model mis-specification, and paralogy problems. These issues can be partially mitigated by employing long segments of DNA (e.g., 20 kb segments of Árnason et al. 2018), but if these regions span one or more recombination breakpoints, they will index multiple phylogenetic histories and violate the assumption of no intralocus recombination that underpins summary coalescent methods (Liu et al. 2009). Likewise, hybrid transfer of genetic material between lineages results in a patchwork of ‘native’ and ‘foreign’ DNA along chromosomes. If 20 kb segments include stretches of both native and foreign DNA, the assumption of a single underlying gene tree for the whole 20 kb would be violated. By contrast, the primary sources of gene tree heterogeneity for retroposons are biological and include ILS and possibly hybridization. This assumes that the presence/absence of retroposon insertions has been carefully scored in genomic screens (Doronina et al. 2019).

Árnason et al. (2018) inferred that extensive hybridization occurred in the evolutionary history of balaenopteroids based on network analysis of 34192 gene trees with PhyloNET (Yu et al. 2014) and D_FOIL_/D statistic (ABBA-BABA) tests on biallelic nucleotide sites (Durand et al. 2011; Pease and Hahn 2015). With these two approaches, they detected as many as seven different gene flow signals (Fig. 6C). Beyond these suggested historical pathways for introgression, there is also compelling genetic and phenotypic evidence for recent hybridization events between blue whales and fin whales (Bérubé and Aguilar 1998), as well as for backcrossing between a female fin whale-blue whale hybrid and a male blue whale (Bérubé and Palsbøll 2018). In the presence of such gene flow, we would not expect the observed distribution of retroposons to be fully consistent with MSC expectations. Lammers et al. (2019) reported a dissonant pattern of CHR2 SINE insertions among balaenopteroids, but did not test whether the conflicting signal is the result of just ILS or instead ILS plus gene flow. Here, we re-analyzed the retroposon data set for Balaenopteroidea (Lammers et al. 2019) to determine whether these data suggest patterns of introgression as inferred by Árnason et al. (2018) using analyses of DNA sequences. We performed the χ^2^ Quartet-Asymmetry Test (Fig. 3) on 35 different quartets of species, and sixteen of 35 tests were statistically significant at p = 0.05 following correction for multiple tests (Table 2).

**Table 2.** Mysticete quartets with statistically significant asymmetry that provides evidence for introgression^a^.

| Quartet of taxa | Split | Obs. # quartets | Exp. # quartets under ILS | $\chi^2$ | p value<sup>b</sup> | Introgression pathway<sup>c</sup> |
| --- | --- | --- | --- | --- | --- | --- |
| RW, MW, GW, SW | **RW MW\|GW SW** | **3210** | | 24.54 | 7.26E-07 | I (MW ↔ SW + BW) |
|  | RW GW\|MW SW | 1749 | 1608.5 |  |  |  |
|  | RW SW\|MW GW | 1468 | 1608.5 |  |  |  |
| RW, MW, GW, BW | **RW MW\|GW BW** | **3684** | | 51.46 | 7.31E-13 | I (MW ↔ SW + BW) |
|  | RW GW\|MW BW | 1813 | 1609.5 |  |  |  |
|  | RW MW\|GW BW | 1406 | 1609.5 |  |  |  |
| RW, MW, FW, SW | **RW MW\|FW SW** | **3055** | | 66.72 | 3.33E-16 | I (MW ↔ SW + BW) |
|  | RW FW\|MW SW | 1853 | 1620.5 |  |  |  |
|  | RW SW\|MW FW | 1388 | 1620.5 |  |  |  |
| RW, MW, FW, BW | **RW MW\|FW BW** | **3480** | | 104.61 | 0.0E-10 | I (MW ↔ SW + BW) |
|  | RW FW\|MW BW | 1965 | 1669.5 |  |  |  |
|  | RW BW\|MW FW | 1374 | 1669.5 |  |  |  |
| RW, MW, HW, SW | **RW MW\|HW SW** | **3168** | | 24.47 | 7.56E-07 | I (MW ↔ SW + BW) |
|  | RW HW\|MW SW | 1683 | 1545.5 |  |  |  |
|  | RW SW\|MW HW | 1408 | 1545.5 |  |  |  |
| RW, MW, HW, BW | **RW MW\|HW BW** | **3578** | | 52.16 | 0.0E-10 | I (MW ↔ SW + BW) |
|  | RW HW\|MW BW | 1742 | 1541.5 |  |  |  |
|  | RW BW\|MW HW | 1341 | 1541.5 |  |  |  |
| RW, GW, FW, HW | **RW GW\|FW HW** | **3318** | | 27.52 | 1.55E-07 | III (GW ↔ HW) |
|  | RW FW\|GW HW | 1695 | 1549 |  |  |  |
|  | RW GW\|FW HW | 1403 | 1549 |  |  |  |
| RW, BW, GW, FW | **RW BW\|GW FW** | **1779** | | 29.09 | 6.89E-08 | IV (GW ↔ BW) |
|  | RW GW\|FW BW | 2000 | 2178 |  |  |  |
|  | RW FW\|GW BW | 2356 | 2178 |  |  |  |
| RW, GW, SW, BW | **RW GW\|SW BW** | **3948** | | 97.85 | 1E-10 | IV (GW ↔ BW) |
|  | RW SW\|GW BW | 1736 | 1468 |  |  |  |
|  | RW BW\|GW SW | 1200 | 1468 |  |  |  |
| MW, GW, SW, BW | **MW GW\|SW BW** | **3726** | | 52.86 | 3.58E-13 | IV (GW ↔ BW) |
|  | MW SW\|GW BW | 1795 | 1590 |  |  |  |
|  | MW BW\|GW SW | 1385 | 1590 |  |  |  |
| MW, FW, SW, BW | **MW FW\|SW BW** | **3754** | | 32.78 | 1.03E-08 | VII (BW ↔ FW + HW); also observed BW ↔ FW in nature |
|  | MW SW\|FW BW | 1651 | 1494.5 |  |  |  |
|  | MW BW\|FW SW | 1338 | 1494.5 |  |  |  |
| MW, HW, SW, BW | **MW HW\|SW BW** | **3666** | | 40.57 | 1.90E-10 | VII (BW ↔ FW + HW) |
|  | MW SW\|HW BW | 1694 | 1518.5 |  |  |  |
|  | MW BW\|HW SW | 1343 | 1518.5 |  |  |  |
| RW, HW, SW, BW | **RW HW\|SW BW** | **3845** | | 83.68 | 1E-10 | VII (BW ↔ FW + HW) |
|  | RW SW\|HW BW | 1598 | 1359.5 |  |  |  |
|  | RW BW\|HW SW | 1121 | 1359.5 |  |  |  |
| RW, FW, SW, BW | **RW FW\|SW BW** | **4108** | | 72.97 | 1E-10 | VII (BW ↔ FW + HW); also observed BW ↔ FW in nature |
|  | RW SW\|FW BW | 1540 | 1320.5 |  |  |  |
|  | RW BW\|FW SW | 1101 | 1320.5 |  |  |  |
| RW, BW, FW, HW | **RW BW\|FW HW** | **3131** | | 32.25 | 1.36E-08 | HW ↔ BW |
|  | RW FW\|HW BW | 1758 | 1597.5 |  |  |  |
|  | RW HW\|FW BW | 1437 | 1597.5 |  |  |  |
| RW, SW,<br>FW, HW | **RW SW\|FW HW** | **3349** |  | 29.50 | 5.60E-08 | HW <-> SW |
|  | RW FW\|HW SW | 1499 | 1357.5 |  |  |  |
|  | RW HW\|FW SW | 1216 | 1357.5 |  |  |  |

<sup>a</sup>The species tree split for each quartet is shown in bold font.
<sup>b</sup>Corrected for multiple comparisons using a Holm-Bonferroni test.
<sup>c</sup>Roman numerals correspond to introgression pathways of Arnason et al. (2018).
Abbreviations: BW, blue whale; FW, fin whale; GW, gray whale; HW, humpback whale; MW, minke whale; RW, North Atlantic right whale; SW, sei whale.

The overall pattern of retroposon insertions is complex and includes multiple quartets of species with skewed support for alternative topologies relative to expectations based on ILS alone (Table 2, Fig. 6D). Even with this complexity there are several themes that emerge from these comparisons. First, minke whale and blue whale are overrepresented in three different quartet tests, as are minke whale and sei whale. These results are consistent with Árnason et al.’s (2018) gene flow pathway I between the minke whale lineage and the common ancestral lineage of blue whale + sei whale (Fig. 6C,D). Second, there is one significant quartet test that is consistent with gene flow between the gray whale lineage and the humpback whale lineage, which corresponds to Árnason et al.’s (2018) gene flow pathway III. Third, there are three asymmetry tests that are consistent with Árnason et al.’s (2018) gene flow pathway IV between the gray whale lineage and the blue whale lineage. Fourth, four comparisons are consistent with Árnason et al.’s (2018) gene flow pathway VII between the common ancestral lineage of humpback whale + fin whale and the blue whale lineage. Two of these four comparisons are also consistent with recently observed hybrids between fin and blue whales. Finally, the remaining two overrepresented quartet splits are (right whale, fin whale|humpback whale, sei whale) and (right whale, fin whale|humpback whale, blue whale) (Table 2). These results are suggestive of hybridization between the humpback whale lineage and the common ancestral lineage of blue whale + sei whale rather than Árnason et al.’s (2018) pathway VI between the common ancestral lineage of fin whale + humpback whale and the common ancestral lineage of blue whale + sei whale.

We note that the interaction of multiple gene flow pathways is expected to result in a complex pattern of retroposon insertions that is not necessarily reflected in the overabundance of observed retroposon insertions for a particular alternative quartet. For example, hybridization between gray whale and humpback whale (path III of Árnason et al. 2018) will increase the proportion of retroposons that are shared between these two species in some quartets, but hybridization between gray whale and blue whale (path IV) and between gray whale and the ancestor of blue whale + sei whale (path V) are expected to have the opposite effect. These competing pathways of gene flow become even more complex if we consider past hybridization between the ancestor(s) of a living taxon and a completely extinct side-lineage of the clade (Maddison 1997; Rogers et al. 2019). Nevertheless, an evolutionary history of hybridization in balaenopteroids is suggested by significant retroposon asymmetry via several pathways that violate expectations based on ILS alone (Table 2; Fig. 6D). Importantly, our comparisons of observed and expected retroposons are not complicated by non-biological sources of gene tree heterogeneity that plague similar comparisons for gene trees that are based on sequence alignments (Song et al. 2012; Zhong et al. 2013; Jarvis et al. 2014; Liu et al. 2017).

## Conclusions

We contend that retroposon insertions provide a superior option to sequence-based gene trees for inferring species trees with summary coalescence methods because retroposons are better matched to the assumptions of these methods, especially for divergences that are closely spaced and deep in the tree of Life (Fig. 5B,D; Springer and Gatesy 2019).First, retroposon insertions are subject to interlocus recombination and ILS, but are not impacted by intralocus recombination. Each retroposon insertion is used to infer just a single, unique bipartition of taxa. Second, gene tree reconstruction error is minimized because retroposon insertions have very low rates of homoplasy. Finally, many or most retroposon insertions occur in genomic safe havens where they are tolerated by natural selection and are effectively neutral.

We analyzed three retroposon data sets (Placentalia, Laurasiatheria, Balaenopteroidea) with ASTRAL and pSVD-Quartets (Fig. 2) to infer species trees that are in general agreement with previous analyses that did not directly account for ILS (Nishihara et al. 2009; Doronina et al. 2017a; Lammers et al. 2019). We also provide three support measures for clades supported by ASTRAL that take ILS into account (Figs. 5–6). Our coalescence analysis of Laurasiatheria retroposons further identified a possible case of the anomaly zone, where the most probable gene tree(s) disagrees with the species tree (Degnan and Rosenberg 2006, 2009) (Fig. 5C,D), for lineage splits that likely occurred in the Cretaceous (Meredith et al. 2011).

As we have argued recently, the analysis of retroposons with ILS-aware methods offers hope for resolving tight sequences of branching events that are deep in evolutionary history (Springer and Gatesy 2019). Here we further demonstrate the utility of retroposon data for discerning ancient hybridization events using a simple but general χ^2^ Quartet-Asymmetry Test (Fig. 3). As for traditional ABBA-BABA tests on DNA sequence data, significance suggests that ILS alone is insufficient for explaining highly skewed distributions of retroposons for particular species quartets. When observed and expected distributions of retroposons for alternative quartets significantly conflict, a plausible explanation for this misfit is introgression because other sources of gene tree heterogeneity such as gene tree reconstruction error and intralocus recombination are thought to be inconsequential for retroposon insertions when these characters are conservatively coded (Doronina et al. 2019). More specifically, variables such as strong rate heterogeneity and heterogeneous base composition can drive asymmetry in the alternative quartets that are inferred from sequence-based gene trees, (Pease et al. 2018), but retroposon insertions are immune to these problems.

In the case of balaenopteroid baleen whales, comparisons between observed and expected distributions suggest that ILS alone cannot account for the observed distribution of retroposons. By contrast, retroposon insertions for Laurasiatheria and for Placentalia are consistent with ILS expectations. We suggest that our general test for asymmetry in retroposon patterns, which allows for all possible outcomes of ILS, offers a valuable framework for the analysis of ancient hybridization events across the Tree of Life in lineages where retroposons were active. The development of novel methods that simultaneously consider both ILS and introgression should be a priority in future systematic studies (Yu et al. 2014; Solís-Lemus et al. 2016). Methods that allow just ILS may recover an incorrect species tree in the face of introgression, even if gene tree inaccuracy is only a minor issue as for retroposon insertions. If there is evidence for introgression, then retroposons may be especially useful for teasing apart different regions of the genome with branching versus reticulate histories.

